# Admixture dynamics in colonial Mexico and the genetic legacy of the Manila Galleon

**DOI:** 10.1101/2021.10.09.463780

**Authors:** Juan Esteban Rodríguez-Rodríguez, Alexander G. Ioannidis, Erika Landa-Chavarría, Javier Blanco-Portillo, Consuelo D. Quinto-Cortés, Rosenda I. Peñaloza-Espinosa, Karla Sandoval, Andrés Moreno-Estrada

**Author notes:** These authors contributed equally (Juan Esteban Rodríguez-Rodríguez, Alexander G. Ioannidis).

## Abstract

Mexico has considerable population substructure due to pre-Columbian diversity and subsequent variation in admixture levels from trans-oceanic migrations, primarily from Europe and Africa, but also, to a lesser extent, from Asia. Detailed analyses exploring sub-continental structure remain limited and post-Columbian demographic dynamics within Mexico have not been inferred with genomic data. We analyze the distribution of ancestry tracts to infer the timing and number of pulses of admixture in ten regions across Mexico, observing older admixture timings in the first colonial cities and more recent timings moving outward into southern and southeastern Mexico. We characterize the specific origin of the heterogeneous Native American ancestry in Mexico: a widespread western-central Native Mesoamerican component in northern Aridoamerican states and a central-eastern Nahua contribution in Guerrero (southern Mexico) and Veracruz to its north. Yucatan shows lowland Mayan ancestry, while Sonora exhibits a unique northwestern native Mexican ancestry matching no sampled reference, each consistent with localized indigenous cultures. Finally, in Acapulco, Guerrero a notable proportion of East Asian ancestry was observed, an understudied heritage in Mexico. We identified the source of this ancestry within Southeast Asia—specifically western Indonesian and non-Negrito Filipino—and dated its arrival to approximately thirteen generations ago (1620 CE). This points to a genetic legacy from the 17^th^ century Manila Galleon trade between the colonial Spanish Philippines and the Pacific port of Acapulco in Spanish Mexico. Although this piece of the colonial Spanish trade route from China to Europe appears in historical records, it has been largely ignored as a source of genetic ancestry in Mexico, neglected due to slavery, assimilation as “Indios” and incomplete historical records.

## 1. Introduction

Population genetics studies have shed light on past historical events and demographic dynamics that in turn are associated with cultural influences [1,2]. Before European contact, the territory of modern day Mexico was occupied by indigenous peoples (Native Americans), who arrived via one or more founding populations that crossed from Asia to the Americas over what is now the Bering Strait ~20,000 years ago [3]. However, these populations exhibit genetic substructure due to their subsequent separation and founder effects. When the Spanish arrived in the Americas in 1492 CE, Mexico did not exist as a unified nation or even as a homogenous culture. Instead, modern Mexico can be broadly classified into two regions based on the lifestyle of its pre-contact Native Americans: Mesoamerica and Aridoamerica. Mesoamerican Natives were characterized by their sedentarism and cultural roots, emerging from a common mother culture: the Olmecs. Mesoamerica extends from western and central Mexico to Central America, comprising several ancient civilizations and ethnic groups. Meanwhile, Aridoamerica consists of the dry region north of Mesoamerica: present-day northern Mexico. Early Aridoamerican natives consisted of ethnically heterogeneous groups practicing nomadic and semi-nomadic lifestyles due to the harsh climate. Upon the Spanish arrival to the Americas, a great part of Mesoamerica, specifically central Mexico, was under the rule of the Mexica (also known as Aztecs) a Nahuatl speaking confederation of city-states [4]. Due to the extent of this Nahuatl speaking empire and the language’s further use by the Spanish as an indigenous lingua franca [5], Nahuatl remained the predominant native language of Mexican territory to the present-day. Nahuatl speakers are referred to as Nahua people. Many other ethnic groups have persisted in the Mexican territory since that time, some surviving under Mexica rule and others resisting subjugation by them [4]. Central Mexican natives such as the non-Nahua Tarascans to the west and the Nahua Tlaxcaltec people remained independent of the Mexica. The latter are known for allying with the Spanish against the Aztecs [4]. The southern Mexican civilization of Tututepec led by the Zapotec and Mixtec, among others, also remained independent [6], as did several Mayan speaking states in the southeast, comprising present-day Chiapas, Yucatan, and parts of Belize, Honduras and Guatemala [7].

The Spanish first conquered the capital of the Mexica Empire, Tenochtitlan now Mexico City, in 1521 CE [8]. Other empires fell shortly after with the help of Spain’s native allies. The Tarascan Empire, located in western Mexico (Michoacan and Jalisco) and ruled by the Purepecha people, lost its independence in 1530 CE. The Purepecha also had an important presence in the Bajio region, comprising Guanajuato, Eastern Jalisco, Aguascalientes and Southern Zacatecas, during the colonial period. In this region, peace was purchased by the Spanish after having lost the Chichimec War in 1590 CE [9]. In this case, semi-nomads were assimilated once they adopted a new sedentary lifestyle, which the Spanish encouraged. In Oaxaca, ethnic groups such as the Zapotec and Mixe persisted culturally. In the southeast territories of the Maya, conquest came in 1543 CE with the entry of Spain into the Yucatan. Mayan presence and culture resistance were nevertheless considerable and long-lived, as reflected in the Caste War of 1847 CE [10]. Vast areas in northwest Mexico also had a long indigenous persistence with very late contact and many failed conquest attempts. Indeed, in Sonora, the first cities were not built until 1700 CE and the area achieved stable settlement only after 1787 CE due to warfare.

Soon after European contact, the territory of what is now Mexico experienced extensive continental admixture. Europeans and sub-Saharan Africans admixed with local indigenous groups, for instance, in the mines of Guanajuato [11]. Around the 17^th^ century admixture began to increase. By the independence of Mexico in the 19^th^ century admixed citizens accounted for up to 40% of the population. Today only 6% of Mexicans speak an indigenous language [12], while most Mexicans speak Spanish and consider themselves “Mestizo,” previously an admixed caste name that is nowadays used with the broad meaning of admixed. Indeed, genetic studies from non-indigenous Mexicans have shown that most such individuals exhibit some degree of admixture involving all three major colonial-era ancestry components (Native American, European, and sub-Saharan African ancestry) [13,14].

Epidemics, droughts, famine, and forced labor caused a collapse of the indigenous economic systems, resulting in a demographic disaster that had its most critical point at 1646 CE. The population of New Spain at its lowest reached ~1,700,000 people [15] with collapse affecting the largest sector at the time: Native Americans. The importation of African slaves was promoted around this period to compensate for the loss of native labor. Population numbers began to increase even though many droughts and epidemics continued over the next century [11]. Some of the most cosmopolitan cities began an unprecedented admixture process, as mining wealth attracted Spanish people and required Native Americans and Africans for labor. Before the collapse, the number of admixed people recorded in 1570 CE constituted only 0.5% of New Spain’s population. By 1810 CE that number reached 39.5% [15]. Nowadays, most Mexicans self-identify as “Mestizo”, making Native Americans a minority in Mexico.

A generalized Iberian source for the European ancestry of Latin American admixed peoples, including Mexico, has been observed. On the other hand, sub-continental structure has been found in the Native American component across Latin American countries with admixed individuals resembling nearby indigenous cultures [16]. However, to date admixture dynamics have not been characterized by region within Mexico, and substructure within the Native American component has not been characterized in high-resolution.

In addition, although Native Americans, Spanish Europeans and sub-Saharan Africans had the largest presence in Mexico during the colony, other ethnic groups immigrated to colonial Mexico, in particular from Asia. These arrived via the Manila Galleons, ships that conducted the trans-Pacific trade with the Philippines every year between 1565 CE and 1815 CE [17]. The largest period of such migration occurred in the 17th century, compensating for the diminished labor force following the indigenous demographic collapse [17]. Some Asians travelled freely to Mexico, but many others were slaves from Manila, where a third of the population were slaves belonging to diverse indigenous groups [17]. The main disembarkation point was in southern Mexico in the Pacific Coastal port of Acapulco, Guerrero. This ancestral contribution has often been overlooked, since Asians were treated as indigenous vassals by law in the 17th century. That is, they were referred to as “Indios” just as Native Americans, and they were assimilated thus into the population [17]. Historical records estimate a total of 40,000-120,000 immigrants from Manila in colonial Mexico [18], and the Spanish wrote they were very numerous in Acapulco, where every Spanish home had at least three, and up to eighteen, Asian slaves [17]. The genetic legacy of this trans-Pacific trade has not been previously characterized in Mexican genomes.

## Results

### 2. Beyond the 3-way admixture model in Mexico: East Asian ancestry

Global ancestry proportions were estimated with unsupervised Admixture [19] including five continental reference populations and admixed cosmopolitan Mexicans from ten sampled cities in ten different Mexican states. Admixture at K=5 distinguished the broad continental components: sub-Saharan African, European, Native American, East Asian and Melanesian (**SI Figure 1**). At this resolution, all pre-contact populations from the Americas are clustered together as Native American (excluding more recent migrations from Asia such as Na-Dene and Inuit) [20]. Moreover, the widespread Austronesian ancestry in Southeast Asia appears as East Asian, the closest reference population to this component.

Ancestry clusters in admixed Mexicans consist of mainly Native American and European, followed in decreasing order by sub-Saharan African, East Asian and Melanesian. These proportions differ within the Mexican subregions as reported in previous studies [13]. For instance, European ancestry is more prevalent in cosmopolitan samples from northern Mexico, especially in Sonora (61.9% in average). Native American ancestry shows higher proportions in southern Mexico, with the highest contribution in Oaxaca (81.9% in average), according to our study. Sub-Saharan African ancestry reaches up to 32.3% in individuals from coastal states known for their Afro-Mexican presence [21], namely Veracruz and Guerrero.

In this study, we included the assessment of a fourth continental origin in Mexico: Asian ancestry. East Asian and Melanesian global ancestries are estimated at less than 4% combined in the majority of cosmopolitan Mexicans. (These two combined genetic components encompass any East Asian, Southeast Asian and Oceanian contributions [22].) Small proportions are not reliable, as they could be Native American ancestry misassigned as East Asian due to both populations sharing a more recent common ancestor to the other references [23,24]. However, some individuals in the dataset exhibit more than 5% of East Asian and Melanesian global ancestry, for instance 12 out of the 50 individuals in the Pacific coastal city of Acapulco, Guerrero, where one individual reached up to 14.5% of East Asian ancestry (**SI Figure 1**). The high proportions of Asian-derived ancestry in these individuals can be attributed to Asian immigration following European contact and not to misassigned Native American ancestry. Moreover, three individuals from Sonora, Oaxaca, and Yucatan showed a combined East Asian and Melanesian ancestry greater than 5%. These admixed Mexican individuals with more than 5% of Asian component in the autosomes provided long enough haplotypes to characterize their within-continent origins across Asia.

### 3. Admixture timing dynamics differ within Mexico

Phasing (separation of haplotypes) and local ancestry were applied to the genotype data using SHAPEIT2 [25] and RFMix v1.5.4 [26], respectively. Local ancestry was estimated using the three most common ancestries in the dataset as references: Native American, European and Sub-Saharan African. Meanwhile individuals having >5% combined East Asian and Melanesian ancestries were excluded from these analyses, in order to avoid of East Asian haplotype classifiers and thus enabling a simple K=3 model. Demographic inference analyses used the block length distribution of local ancestry assigned genomes withing each population via the Tracts algorithm [27]. This method allows testing for multiple migration waves from the same source population, and offers a strong approach to this dataset, as it is not sensitive to ascertainment bias and as it allows more complex demographic models to be tested, in contrast to LD-based methods such as MALDER [28]. However, it cannot handle more than three ancestries, forcing the demographic inference to be limited to the three most common ancestries previously mentioned.

We tested four Tracts models for each cosmopolitan Mexican population, generating a likelihood score from the adjustment between the model and real data (**SI Table 1**). The models with the best likelihood were chosen for each population, resulting in contrasting admixture dynamics and timings (**Figure 1**). All models found an initial admixture event between Native Americans and Europeans followed by an African pulse. This dynamic coincides with historical data, as the largest African slave influx in colonial Mexico took place decades after the main contact between Europeans and Native Americans [29]. Two contrasting ranges of initial timings are observed in the dataset. On the one hand, states in the Southeast and adjacent areas such as Oaxaca in the South and Veracruz in the Gulf exhibit more recent admixture timings between 9 and 11 generations in the past (1680 CE–1740 CE), regions with a higher Native American presence and ancestry proportion in their cosmopolitan populations [13]. The rest of the country showed earlier estimates, between 14 and 16 generations in the past (1530 CE–1590 CE). The latter shows that, in some regions of Mexico, admixture occurred within the first century of European contact, while in others it occurred in the 17^th^ century, after the enormous reduction of the natives at the same time that the admixed caste growth became considerable [15].

**Figure 1.**
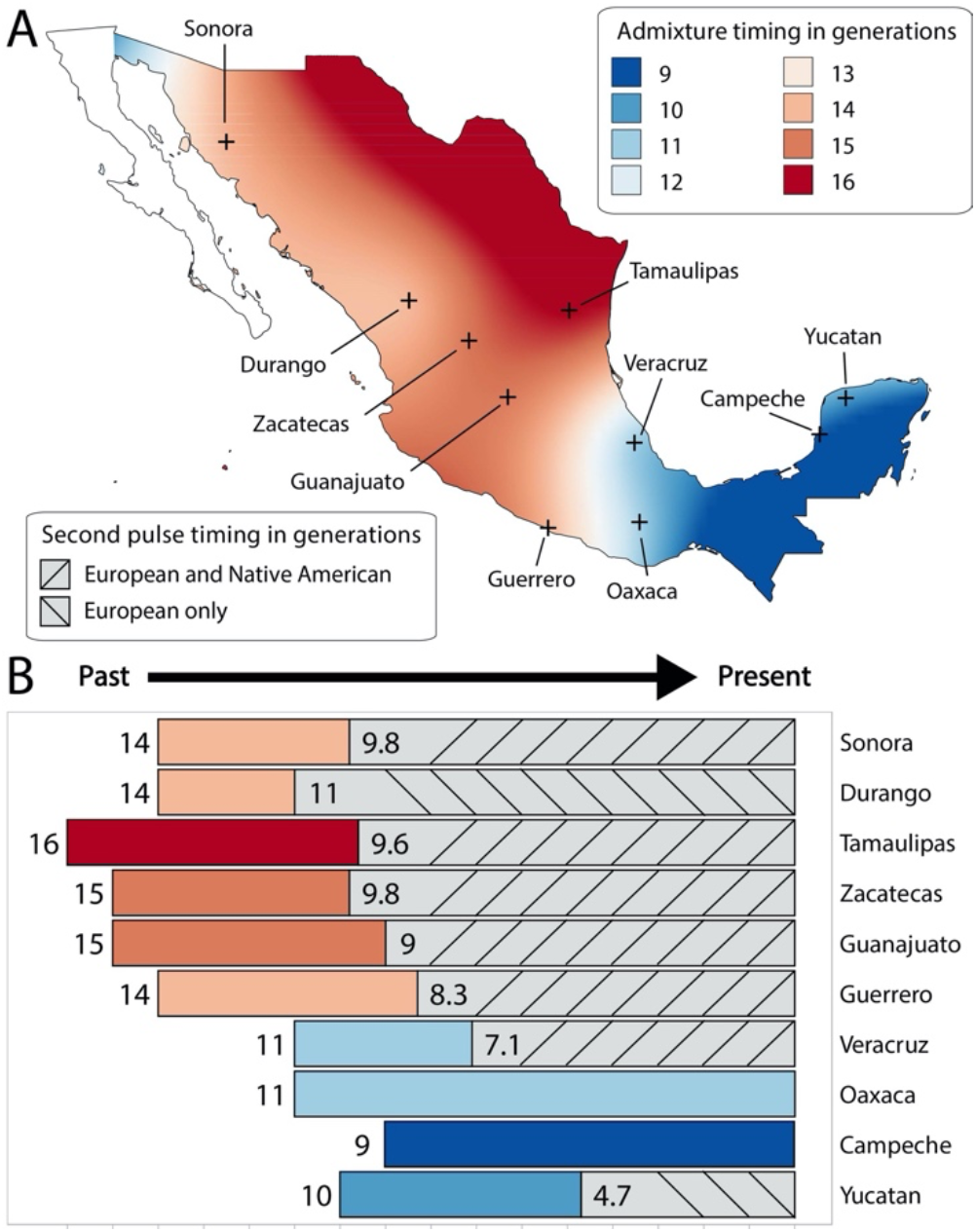
Admixture timings across cosmopolitan populations by Mexican state. **A)** The density map shows the inferred date of the first admixture event predicted by Tracts for each sampling location. Intermediate space between data points is filled with interpolated values in MapViewer according to the observed adjacent estimates. Values indicate how many generations in the past the admixture event between Native Americans and Europeans resulted in the initial admixed population. Crosses represent the sampling locations within the state, namely Hermosillo (Sonora), Ciudad Victoria (Tamaulipas), Zacatecas City (Zacatecas), Guanajuato City (Guanajuato), Acapulco (Guerrero), Xalapa (Veracruz), Oaxaca City (Oaxaca), Campeche City (Campeche) and Merida (Yucatan). The Baja California Peninsula has been excluded as no sampling location were present on the peninsula. Caution should be used to draw conclusions from areas with no data, as well as intermediate locations between sampling points. **B)** The bar plot shows the demographic model that best fits the data per state, including the inferred timing of migration pulses (indicated by number of generations in the past) and the type of each model. Older timings are shown on the left while recent events are on the right side. The solid section of the bars represents the initial admixture event between Native Americans and Europeans, while the remainder of the bars with patterned lines represent the occurrence of further incoming migration pulses into the already admixed population. In most cases these represent a dual pulse of Native American and European ancestry into an already admixed population. Only Durango and Yucatan show a second pulse of solely European ancestry. Oaxaca and Campeche did not exhibit second pulses after the initial admixture event.

### 4. Highly sub-structured Native American ancestry in cosmopolitan Mexicans

Native American substructure across indigenous and cosmopolitan populations was analyzed with ancestry specific methods: Multiple Array Ancestry Specific Multidimensional Scaling (MAAS-MDS) [30], an MDS designed for analyzing samples from several different genotyping arrays simultaneously. Usually, genetic analyses rely on the intersection of all sets of genetic markers, resulting in extremely limited numbers of markers remaining to perform any kind of inference. By overcoming this technical difficulty, we can include more populations by combining five different published genotyping array datasets (details about these runs are provided in **Table 1**).

**Table 1.**

| Analysis details | Target populations | Individuals | Markers |
| --- | --- | --- | --- |
| **Admixture** |  |  |  |
| Array A and B<br>(Illumina 550K) | **Cosmopolitan Mexican <sup>1</sup>:**<br>Sonora, Durango, Tamaulipas, Zacatecas,<br>Guanajuato, Veracruz, Guerrero, Oaxaca,<br>Campeche and Yucatan | 369 | 509,426 |
| **Tracts** |  |  |  |
| K3 Tracts timings | **Cosmopolitan Mexican <sup>1</sup>:**<br>Sonora, Durango, Tamaulipas, Zacatecas,<br>Guanajuato, Veracruz, Guerrero, Oaxaca,<br>Campeche and Yucatan | 354 | 803,636 |
| K4 Asian Tracts timings | **Cosmopolitan Mexican <sup>1</sup>:**<br>Guerrero* | 11* | 803,636 |
| **Native American MAAS-MDS** |  |  |  |
| **RFMix run**<br>Array A<br>(Affymetrix 500 K and<br>Illumina 550 K) | **Cosmopolitan Mexican <sup>1</sup>:**<br>Sonora, Tamaulipas, Zacatecas,<br>Guanajuato, Veracruz, Guerrero and<br>Yucatan<br>**Native American <sup>1</sup>:**<br>Tepehuan, Northern Zapotec and Maya<br>(Campeche) | 310 and 70<br>Total: 380 | 789,054 |
| **RFMix run**<br>Array B<br>(Illumina 550K) | **Cosmopolitan Mexican <sup>1</sup>:**<br>Durango, Oaxaca and Campeche | 57 | 518,410 |
| **RFMix run**<br>Array C<br>(Affymetrix 6.0) | **Native American <sup>2</sup>:**<br>Tarahumara, Huichol, Purepecha, Nahua<br>(Jalisco), Nahua (Puebla), Totonac,<br>Mazatec, Northern Zapotec, Triqui, Tzotzil<br>and Maya (Quintana Roo) | 242 | 794,029 |
| **RFMix run**<br>Array D<br>(Axiom LAT 1) | **Native American <sup>3</sup>:**<br>Nahua (Mexico City), Nahua (Veracruz)<br>and Nahua (Guerrero) | 48 | 722,489 |
| **RFMix run**<br>Array E<br>(Illumina 610-Quad and<br>Illumina 650Y) | **Native American <sup>4</sup>:**<br>Pima, Yaqui, Purepecha, Mixtec, Southern<br>Zapotec, Mixe and Maya (Quintana Roo) | 94 | 364,396 |
| **East Asian MAAS-MDS** |  |  |  |
| **RFMix run**<br>Array A<br>(Affymetrix 500 K and<br>Illumina 550 K) | **Cosmopolitan Mexican <sup>1</sup>:**<br>Sonora*, Guerrero* and Yucatan* | 14* | 803,636 |
| **RFMix run**<br>Array B<br>(Illumina<br>HumanHap550K) | **Cosmopolitan Mexican <sup>1</sup>:**<br>Oaxaca* | 1* | 518,409 |
| **RFMix run**<br>Array C<br>(Affymetrix 6.0) | **East and Mainland Southeast Asia <sup>5</sup>:**<br>Japan, Northern China, Southern China,<br>Vietnam<br>**Philippines <sup>5</sup>:**<br>Mindanao and Negrito from Mindanao<br>**Indonesia <sup>5</sup>:**<br>Sumatra, Borneo, Lesser Sunda Islands and<br>Maluku Islands<br>**Oceania <sup>5</sup>:**<br>Fiji and Papua New Guinea | 225 | 561,339 |
| **RFMix run**<br>Array D<br>(Illumina OmniExpress<br>Bead Chips and whole<br>genome) | **Philippines <sup>6</sup>:**<br>Igorot, Luzon and Visayas | 22 | 542,878 |

\* Only individuals with >5% East Asian + Melanesian local ancestry were included in the respective analysis.
1. Mexican Genome Diversity Project (MGDP) from [13]. 2. Native Mexican Diversity Project (NMDP) from [13]. 3. Nahua populations from [42]. 4. Native American reference panel from [49], only populations from Mexico were included. 5. Southeast Asian reference panel from [50]. 6. Filipino populations from Southeast Asian reference panel [47] and whole genome Igorot from Simons Genome Diversity Project [48]

Native American ancestry in cosmopolitan Mexicans shows a considerable substructure, these differences match geography to some extent, as previously observed [13]. However, in this study we elucidated a precise origin for these haplotypes matching modern Native American groups in Mexico. The most distant cosmopolitan populations from central Mexico exhibited more differentiated Native heritages. For instance, the northwestern border state of Sonora had some affinity to Aridoamerican peoples such as the Yaqui, Tarahumara and Tepehuan, while the southeastern states of Campeche and Yucatan clustered predominantly with the local Maya from the peninsula (**Figure 2**). Most of the states clustered with western, central and eastern natives. Individuals from Durango, Tamaulipas, Zacatecas and Guanajuato exhibited an intermediate affinity with western and central natives, the former including the Purepecha and Nahua from Jalisco and the latter consisting of the Nahua from Mexico City and Guerrero. Populations from Veracruz and Guerrero did not cluster with western natives, instead they exhibited central and eastern native ancestry, i.e. Nahua from Mexico City, Guerrero, Veracruz and Puebla, as well as Totonac. The Nahua from Puebla represent the geographically nearest Nahua population to the sampled city of Xalapa, Veracruz. Finally, Oaxaca exhibited a unique ancestry compared to the other cosmopolitan Mexicans, showing Mazatec, Zapotec, and Mixtec affinities. These Native American groups have had an important presence in this state since pre-contact times. The admixed samples from the Valley of Oaxaca overlap with Zapotecs from the Sierra Sur region and Mixtecs from the Valley of Oaxaca, in contrast to other Zapotec populations from the Sierra Norte region and the Triqui people from the Mixteca region (western Oaxaca).

**Figure 2.**
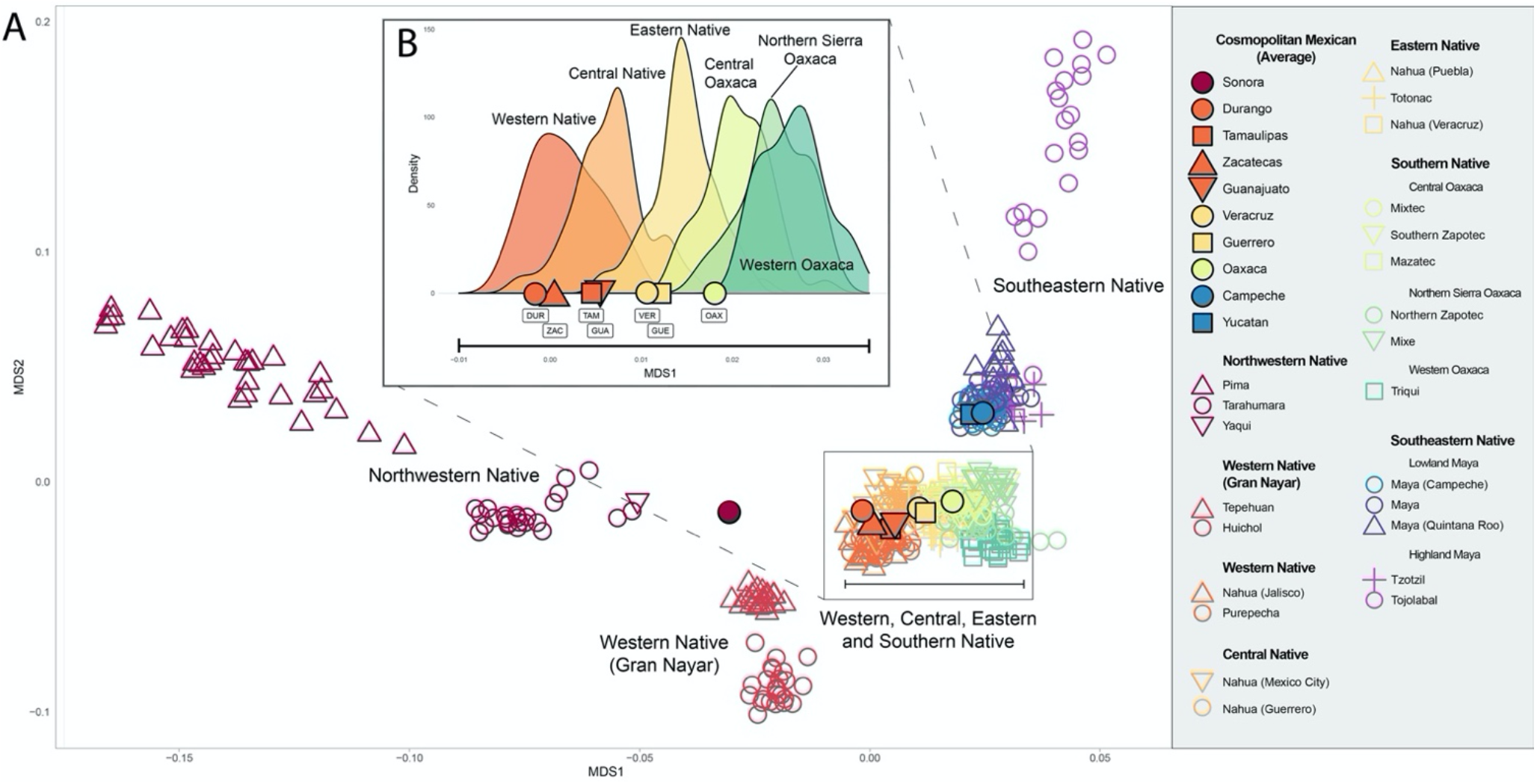
Native American substructure in cosmopolitan Mexican states. **A)** Native American individuals are shown with empty figures, and cosmopolitan Mexican populations are shown as an average per state with filled shapes. Both Native Americans and cosmopolitan Mexicans have their non-native ancestries masked. Individuals with <10% of Native American ancestry were not included. **B)** Populations exhibiting western, central, eastern and southern native affinities were included in the inset figure as shown with the rectangle. MDS 1 corresponds to the x-axis in the density kernel plot. Native American populations were grouped into geographic and genetic categories and plotted as densities, while cosmopolitan Mexican averages were plotted as single points on the x-axis according to their MDS 1 projection.

The affinity between cosmopolitan Mexicans and nearby modern Native Americans supports a clear genetic continuity in places such as Sonora, Veracruz, Guerrero, Oaxaca, Campeche, and Yucatan. Sonora, in the Northwest of Mexico, shows affinity to some extent with northwestern natives; Veracruz and Guerrero cluster with close Nahua samples from central and eastern Mexico; Oaxaca overlaps with local Oaxaca natives, and Campeche and Yucatan resemble the Maya from the Yucatan Peninsula. On the other hand, cosmopolitan Mexicans from the near north, northeast and north-central states, while located in Aridoamerica, show Mesoamerican affinity. However, caution should be taken as the Aridoamerican populations from these regions are not sampled and most are no longer extant; thus, we cannot discard a genetic contribution from these pre-contact hunter-gatherers.

### 5. Native American genetic substructure and linguistic affinity

Nahua peoples (or Nahuatl speakers) have been the most numerous natives in Mexico since European contact. Nahuatl was the language of the extensive Mexica Empire, Tlaxcaltec peoples (allied to the Spanish) and other previous civilizations in the Post-Classic period, such as the Toltecs. Moreover, upon the fall of the Mexica capital, Tenochtitlan, the Spanish rulers promoted the use of the Nahuatl language. This ethnic category is based in a shared language family group that extends from northern Mexico, such as the Mexicanero peoples in Durango, to the country of El Salvador, where the Pipil language is spoken. However, do all these populations share a common genetic profile the same way they share their language? Previous studies have shown Nahua peoples do not make a monophyletic clade, instead they appear in several branches of Mesoamerican natives from Mexico with interspersed, unrelated linguistic families [13]. To explore this further, we included additional Nahua populations in this study, and we identified three main groups of Nahua peoples that overlap or resemble neighboring non-Nahua natives.

The most differentiated Nahua population is in western Mexico, specifically in the state of Jalisco. This sampling location corresponds to the western Mesoamerican cultural region, where the neighboring Purepecha peoples are located and shows a close affinity with them. A less differentiated genetic profile is seen between central, eastern and southern Mexican natives. Their averages differ, but they overlap slightly with the neighboring regions. The central cluster includes the Nahua from Mexico City and central Guerrero, which exhibit a similar variance. The eastern cluster includes the Nahua from Puebla and central Veracruz, as well as the nearby Totonacs. Finally, southern natives include the heterogeneous natives from Oaxaca. The closest to the eastern Natives are the Mazatec, Mixtec and Zapotec, while the Mixe and Triqui have a particular differentiation.

Linguistic hypotheses regarding diversification and migration patterns from Nahua populations agree with the genetic profiles observed in the MAAS-MDS. Western Nahua are closer to Huichol and Tepehuan populations from the Gran Nayar region in comparison to central and eastern Nahua. Linguistic mutations specific to the western Nahua clade have been proposed to be caused by an interaction with neighboring Corachol groups, which include the Huichol people. Another agreement is observed with the genetic affinity of a central Nahuatl speaking population as intermediate between western and eastern Nahuatl populations. According to linguistic data, it has been proposed that central Nahuatl dialects could have been a mixture of local eastern Nahuas and incoming western Nahuas, after the latter already had interacted with the Cora and Huichol peoples [31]. However, these interpretations should be taken with caution, as the sampling of Nahua and neighboring natives is still poor. More conclusive results can be obtained with the use of more elaborate bioinformatics tools, and with the genotyping of more populations from each Nahua clade, as well as from the equally underrepresented neighboring natives, with whom the Nahua could have admixed.

Aside from the Nahua, other groups' affinities can be noted. Some natives from Oaxaca show similarities across languages, possibly reflecting interethnic interactions, gene flow or language shifts. For instance, the Zapotecs from the Sierra Sur region overlap better with the Mixtec and Mazatec peoples, compared to other Zapotecs from the Sierra Norte region, who appear somewhat closer to the neighboring Mixe. More sampling locations are required to make inferences from this, as Mixtecs extend over a vast area and could have significant genetic substructure.

### 6. Heterogeneous origins of the Asian ancestry in Mexico

In order to pinpoint the origin of the Asian component in Mexico, a MAAS-MDS was performed with a reference panel of East Asian, Southeast Asian and Oceanian populations. Cosmopolitan Mexicans having more than 5% combined East Asian and Melanesian ancestry were included, resulting in one individual from Sonora, one from Oaxaca, one from Yucatan and twelve from Guerrero. Sonora and Yucatan grouped near Chinese reference populations; Oaxaca clustered broadly with maritime Southeast Asia; while Guerrero showed a heterogeneous profile (**Figure 3**). No cosmopolitan Mexican sample showed Melanesian variation; therefore, the MAAS-MDS plot was zoomed into, excluding populations with Melanesian contributions, for visibility (complete MDS plot is on supplementary information **SI Figure 2**). Most individuals from Guerrero clustered with maritime Southeast Asia, except for one individual positioned near southern China. Individuals from Guerrero resemble western Indonesian and non-Negrito Filipino populations, specifically those from Sumatra, Mindanao, Visayas and Luzon. Admixture dating of these Asian haplotypes in Guerrero using Tracts fit a single pulse admixture model at 13 generations ago, or in 1620 CE using 30 years per generation (**Figure 4**).

**Figure 3.**
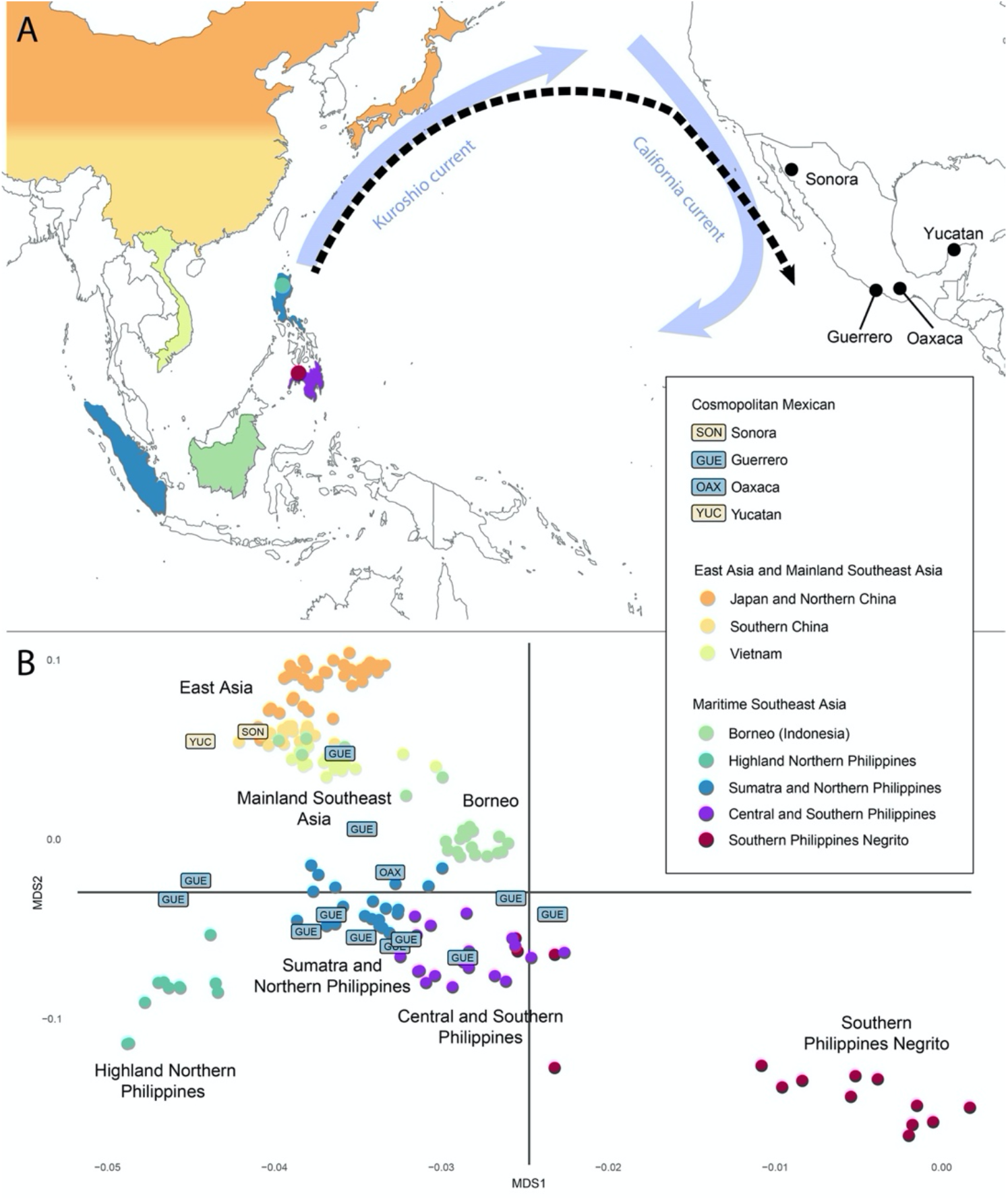
East and Southeast Asian substructure in cosmopolitan Mexican individuals. **A)** Map shows the sampling locations from East and Southeast Asian populations included in the Asian MDS. Sampling locations shown in the map as circles represent isolated populations with contrasting genetic profiles. Cosmopolitan Mexican populations with individuals exhibiting >5% Asian ancestry are shown in the map with black points, most individuals were sampled in Guerrero. The Manila Galleon passage from the Philippines to Acapulco, Guerrero is shown approximately on the map with a black dashed arrow. Blue arrows show ocean currents exploited for this eastward trip. The Pacific Ocean extent is not shown to scale. **B)** An MDS shows the East and Southeast Asian reference individuals with filled circles, while cosmopolitan Mexican individuals are plotted with rectangular labels. The color code in the reference panel coincides with the sampling location on the map, while cosmopolitan Mexican label colors approximately match the native population with which they have the most affinity.

**Figure 4.**
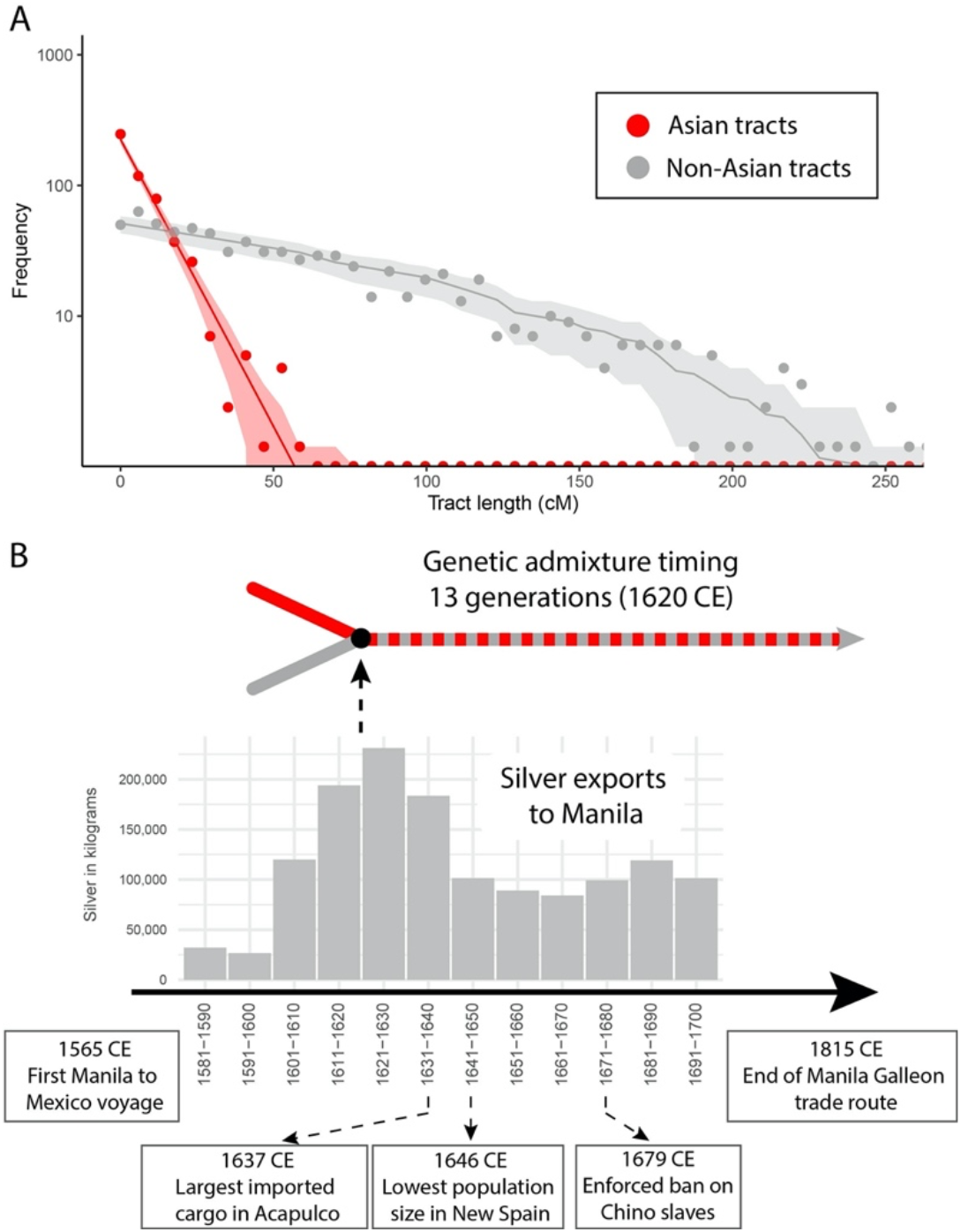
Southeast Asian admixture timing in Acapulco, Guerrero, and Manila Galleon trade data. **A)** Histogram shows the frequency of Asian and Non-Asian derived tracts by length. The expected histogram corresponding to the Tracts model is represented with a solid line and a shaded confidence interval, while the empirical tract length data is shown with points. Asian tract lengths represent East Asian and Melanesian merged ancestries. Non-Asian tracts consist of merged African, European, and Native American ancestries. **B)** Admixture between Asian and Non-Asian ancestry is represented with an arrow diagram, estimated thirteen generations ago by a Tracts analysis. Historical information is provided as a comparison to genetic estimates. In particular, during the Manila Galleon trade period silver exports from Acapulco to Manila are shown with a bar plot as a proxy for voyaging and trade volume, including slave importation. The largest registered cargo from Manila to Acapulco and the final enforced ban on Chino slaves indicated.

This coincides with the Manila Galleon slave trade during the colony, which had a period of activity from 1565 to 1679 CE [17]. This slave trade route originated after the need for additional labor arose due to the demographic collapse of the native populations, and ended when these Asian slaves, mostly residing in Spanish colonial Asia, were actively declared indigenous vassals of the crown and thus free. At that time the Atlantic slave trade from Africa became predominant over the Pacific route [17]. This Southeast Asian component from the Manila Galleon trade could have extended to neighboring coastal Pacific areas of southern Mexico, as could be the case of the individual from Oaxaca. Moreover, although historical records report the residence of “Chinos” predominantly in Guerrero, smaller numbers are also recorded in places such as Colima, Guadalajara, Zacatecas, San Luis Potosi, Veracruz, Puebla, Toluca and, in particular, Mexico City [17]. Thus, we do not rule out the presence of this component in the other populations from the study due to insufficient sampling or statistical power, as well as locations not considered in this study.

On the other hand, East Asian ancestry in Sonora and Yucatan, both distant locations from Guerrero, could possibly represent post-colonial migration events, such as Chinese immigration, mainly from the Guangdong Province, into northern Mexico [32] and the immigration of Korean henequen workers into the Yucatan Peninsula, both occurring during and after the Porfiriato Period (between 1880 and 1910 CE) [33]. However, more extensive sampling across the country is needed to shed light on these genetic signals in order to associate them with these post-colonial historical events.

## 7. Discussion

### Mesoamerican ancestry in Northern Mexico: the pacification of the north

Northern Mexico was difficult for the Spanish to occupy when they first arrived in the Americas due to the arid climate, long distances, and bellicosity of the hunter-gatherer natives. The conquest of the north was not achieved until the center and south were completely pacified, and only after the establishment of a stable economy based on the mining of silver and gold [11]. Northern Mexico, that is the region north of Mesoamerica, was scarcely populated. The pacification and settlement of the northern lands were carried out with the help of Mesoamerican native allies, e.g. Otomi, Mazahua, and Purepecha along with Nahua from the Valley of Mexico, such as the Mexica and Tlaxcaltec [15]. The migration of Mesoamerican natives promoted sedentary lifestyles in Aridoamerica and could explain the native ancestries of admixed northern Mexicans. The Native American ancestry of cosmopolitan samples from Durango, Tamaulipas, Zacatecas and Guanajuato clusters with Mesoamerican ancestries: the Purepecha and Nahua from Jalisco and Mexico City in the Native MAAS-MDS. Durango city originally was populated by the Tepehuan peoples [34], before they were reduced by the epidemics spread by the missions. In this case, cosmopolitan Mexicans from Durango do not resemble the present day local Aridoamerican natives (Tepehuan individuals), as shown in the Native MAAS-MDS, instead they cluster with the Mesoamerican groups to their south. However, a shared affinity between Mesoamericans and northern hunter-gatherers is not ruled out as a possible explanation, as the pre-contact genetic profile of most Aridoamerican groups remains uncharacterized, especially the so-called Chichimeca peoples. The genotyping or sequencing of further modern and ancient Native American groups will shed light on the matter, as many ethnic groups from northern Mexico have not been analyzed and many disappeared as distinct groups after European contact.

### Mine exploitation and admixture in Guanajuato City and Zacatecas City

Guanajuato City and Zacatecas City had their first significant occupations with the formal exploitation of their mines, mainly silver, in 1564 CE. This industry led to the substantial admixture of peoples from diverse continental origins. This involved European descent peoples, who managed the extracted minerals, as well as sub-Saharan African and Native American individuals that worked the mines. Admixture intensities in Guanajuato city and Zacatecas city were unprecedented at that time and our results provide a signal consistent with the three-way mixing occurring during that period, according to the Tracts models 15 generations ago or 1560 CE (**Figure 2**). At European contact, Guanajuato and Zacatecas were populated by hunter-gatherers such as the Guamare and Zacateco people. However, the settlement of the North, mediated by the Spanish with the help of Mesoamerican allies, probably replaced the genetic signal of the hunter-gatherers. In the case of Guanajuato City, these Mesoamericans worked in the mines, leading to admixture. Historical records suggest these Mesoamerican peoples consisted of two main migration sources: Purepechas brought from the state of Michoacan (the former Tarascan Empire) and natives from central Mexico, the Otomi, Mazahua and Nahua peoples [11]. These two native sources could explain the bimodal affinity of the Native American ancestry in Guanajuato City (**SI Figure 3**). The exploitation of the silver promoted a stable economy that allowed for the further expansion and population growth of these admixed mining populations, which then sent expeditions further north to settle Sonora founding Alamos, another silver mining city, in 1682. From Alamos came funding and settlers to establish Alta California (now the US state of California) beginning with the Presidio of Monterey in 1775. Thus, the legacy of admixture in these early mining cities, whose populations later founded the cities of northernmost Mexico, could explain the equally early admixture timings that we find across these much later settled northern states.

### Cultural influence from local Opatan and Cahitan loanwords in Sonoran Spanish and Mayan loanwords in the Yucatan Peninsula’s Spanish

Ancestry specific analysis shows the differentiated component in the Native American ancestry from Sonora. Even though ethnic groups from the same state and surrounding areas are included in the MDS, they define their own cluster nearby, suggesting a native origin of an unsampled group with some similarities to the groups of northwestern and western Mexico. This unsampled native source could be linked to ethnic groups from Sonora and neighboring areas, for instance extant indigenous northern populations not included in the MDS are, Mayo, Guarijio, Mountain Pima and Apache, as well as several extinct populations such as the Opata peoples and the several natives from the neighboring state of Sinaloa, whose population was decimated early in the colonial period, ultimately disappearing [35]. We hypothesize the most probable source population are the Opata peoples, consisting of the linguistically and culturally related Eudeve, Tehuima and Jova peoples. These were the most numerous Native American groups from Sonora until they disappeared in the middle nineteenth century, probably due to cultural assimilation and intermarrying [36]. They rapidly converted to christianity and frequently intermarried with Spanish, in contrast to the neighboring natives (Pima, Seri and Yaqui included in the MDS), who are known for their cultural persistence and hostility towards the Spanish and Mexican governments. An Opatan or Cahitan (the linguistic family to which Yaqui belongs) origin of the native ancestry in Sonora could explain the linguistic influence from these peoples on the local Spanish dialect. Admixture could explain why Sonoran Spanish makes use of native words from these ethnicities, while they remain absent in other Mexican Spanish dialects. Besides the several Opatan and Cahitan toponyms in Sonora, loanwords from both families are commonly used in the Spanish of northwestern Mexicans that do not necessarily identify as indigenous people. For example, Opatan borrowings such as *catota* (marble), *chigüi* (turkey), *sapeta* (cloth diaper) and *tépari* (Tepary bean, *Phaseolus acutifolius*) are used in Sonora, especially in northeast Sonora, the Opateria region. Cahitan borrowings such as *mochomo* (ant), *buqui* (child) and *bichi* (naked) are also frequently used in northwestern Mexico, but not outside [37]. More comprehensive sampling is needed to clarify the origin of this native ancestry, as many extant and extinct populations are not yet genotyped. Also, we cannot discard a combination of native ancestries such as a northwestern and western native admixture due to northward movements of indigenous or admixed Mexicans.

The Yucatan Peninsula, where the cosmopolitan Mexican populations of Campeche and Yucatan are located, has been consistently populated by Mayan peoples since millennia before European contact [38]. The MDS suggests a genetic continuity in the native component of present-day admixed Mexicans from the region. These cosmopolitan Mexicans cluster with lowland Mayans, in contrast to highland Mayans such as the Tzotzil from Chiapas. The genetic affinity coincides with a large cultural influence from the Maya to the cosmopolitan Mexicans from the Yucatan Peninsula, who do not necessarily self-identify as Maya. The cultural influence includes many toponyms, Mayan surnames and linguistic influence into the local southeastern Mexican Spanish dialect. For instance, the most frequent surnames in the Yucatan Peninsula have a Mayan origin, e.g. Pech, Chan, Canul, May, Chi, etc. [39], in contrast to most regions in Mexico where Spanish surnames are by far the most common. This does not dismiss native ancestry in other regions of Mexico, it only shows a correspondence of Mayan heritage between surnames and autosomal genetic profiles of present-day cosmopolitan Mexicans. Spanish in the Yucatan Peninsula (including the Spanish spoken by non-indigenous identifying Mexicans) has been heavily influenced by Yucatec Maya, resulting in considerable phonetic changes and several loanwords, such as *turix* (dragonfly), *huech* (armadillo), *mulix* (curly hair), *xic* (armpit) and *xix* (crumbs) [40].

### Genetic footprint of Asian immigration through the Manila Galleon

Southeast Asian ancestry was observed in Mexicans from Guerrero, particularly from the Pacific port of Acapulco. This profile suggests a genetic remnant from the Manila Galleon, which used Acapulco as the port of disembarkation in Mexico. Limited historical records indicate that the proximal source of these thousands of “Chinos” was the Philippines. Our genetic results revealed some Filipino ancestry together with ancestry related to Sumatra in modern Indonesia, then under Muslim Malay rule.

Although the Pacific trade occurred between Manila and Acapulco, the heterogeneity of Asian ancestry in Acapulco can be explained by the multiethnicity of Manila as there was an active slave trade across the region between the Portuguese of Malacca, the Spanish, and even the Filipino elites targeting the Muslim ruled southern islands in particular via the colonial-era concept of “just war”. Indeed, Spanish slaves from Sumatra are well documented, for instance Magellan’s Malay-speaking slave Henrique, believed to be the first human to have circumnavigated the globe. During the Spanish-Moro conflict, sources suggest that soldiers enslaved more than 4,000 Muslims between 1599 and 1604 alone. These Muslim Filipinos, named Moro by the Spanish, inhabited the southern Philippines. The genetic affinity of one individual from Guerrero with Mindanao (the southernmost major island in the Philippines) suggests an ancestry originating in this context. Most of these captives were sold in the Manila slave market [17]. The cultural impact of this migration is evident in Mexico with the usage of terms of Filipino etymology such as “parián” [41]. Also, the Filipino beverage “tuba,” a coconut wine, which had an important industry in the Pacific coast of Mexico and is still traditionally produced in the coastal region of Colima. People from the coast of Guerrero still recognize this Asian heritage in the region.

Overall, our results reveal an understudied origin for historically neglected passage of large numbers of Asian immigrants into Mexican territory during the colonial period. These origins suggest that revealing an untold history of the Asian slave trade in Mexico can be pursued through the genetic footprint of present-day admixed populations.

## 8. Methods and Methods

### (a) Cosmopolitan Mexican dataset

In order to study the substructure of admixed Mexicans, a total of 369 cosmopolitan Mexicans from ten sampling locations were analyzed and genotyped as part of a previous publication [13]. We utilized the Mexican Genome Diversity Project (MGDP) dataset, which consists of seven cosmopolitan populations genotyped with two microarrays Affymetrix 500K and Illumina 550K, and three cosmopolitan populations genotyped with one microarray, Illumina 550K. The dataset includes 48 individuals from Hermosillo, Sonora; 17 from Ciudad Victoria, Tamaulipas; 19 from Durango City, Durango; 50 from Zacatecas City, Zacatecas; 48 from Guanajuato City, Guanajuato; 50 from Xalapa, Veracruz; 50 from Acapulco, Guerrero; 18 from Oaxaca City, Oaxaca; 20 from Campeche City, Campeche, and 49 from Merida, Yucatan. The cities are among the largest and most important from each Mexican state sampled. All individuals were asked if their four grandparents were born in the state in which they were sampled, thus describing regional admixture events. All analyses performed throughout the paper are focused on these ten cosmopolitan samples, including the dating of admixture timings, and characterizing Native American and Asian substructure differences between Mexican populations.

### (b) Native American reference panel

Previously genotyped samples from the Native Mexican Diversity Project (NMDP) were used to build a reference panel representing the major ancestry components of indigenous populations throughout Mexico. Sampling locations and genotyping details are described in [13]. Additional samples from Nahua populations previously genotyped in a separate study [42] were incorporated to provide better resolution in resolving the substructure in central Mexico. This included 49 self-identified Nahua individuals from San Pedro Atocpan and Xochimilco in Mexico City, Necoxtla in Veracruz and Zitlala in Guerrero. All individuals were Nahuatl speakers with local ancestors. Samples were collected with informed consent permitting population genetic studies. DNA was extracted from blood samples and genotyped using the Axiom LAT 1 array (World Array IV chip), which includes 783,856 SNPs. Genotyping was performed at the Institute for Human Genetics of UCSF and data originally reported as part of the study by [42].

### (c) Global ancestry with Admixture

The analysis was performed with Admixture version 1.3.0 in unsupervised mode. The proportion of five well-differentiated, continental source populations was determined for all samples: sub-Saharan African, European, Native American, East Asian and Melanesian. Each continental signal was estimated with an equal number of reference samples when possible. In order to include the largest number of markers, the cosmopolitan Mexican dataset was merged with whole genome reference data from the 1000 genomes consortium dataset [43] and the Human Genome Diversity Project (HGDP) [44]. Individuals from 1000 genomes provided four continental reference panels, while HGDP provided additional Native American individuals plus the Melanesian continental references. 65 YRI from 1000 genomes represented the sub-Saharan African panel. 65 IBS from 1000 genomes made the European panel. 27 PEL and 2 MXL from 1000 genomes and 36 HGDP individuals from the Americas with >99% of Native American ancestry made the Native American panel. 33 KHV and 32 CHS from 1000 genomes composed the East Asian panel. Finally, 16 HGDP individuals from the Papua New Guinea highlands with >99% of Australo-Papuan ancestry comprised the Melanesian panel. A total of 509,426 SNPs was considered for the Admixture run.

### (d) Phasing with SHAPEIT2 and continental local ancestry assignment with RFMix

Each continental reference panel and admixed Mexican panel were phased separately with SHAPEIT2 and default parameters. Phased haplotypes were given to RFMix version 1.5.4 [26]. The rephasing step was performed with the PopPhased flag due to the absence of trios and duos in the sample set. Default parameters were used, consisting of 0.2 cM long windows, 8 generations, 100 trees to generate per random forest, zero EM, and one for the minimum number of reference haplotypes per tree node. We considered three or five continental reference panels in the local ancestry pipeline depending on the analysis performed. We used the same individuals from 1000 genomes and HGDP as in the Admixture analysis. For the Tracts demographic inferences, three references were used: sub-Saharan African, European and Native American, as Tracts models with four ancestries are non-existent and East Asian ancestry is not considerable after removing the individuals with >5% combined East Asian and Melanesian ancestry. The rest of the analyses employed an additional East Asian and Melanesian reference panel. Details for each analysis and run are provided in **Table 1**, specifying number of markers considered and the populations included for the local ancestry calls.

### (e) Demographic inferences with Tracts models: three continental ancestries

This analysis included tracts from the three most common ancestries in cosmopolitan Mexicans: sub-Saharan African, European and Native American. The few individuals with any Asian ancestry were excluded from this analysis to avoid a change in the distribution of the ancestry tracts that would necessitate modeling even more complex combinations of pulses. The fit of the predicted and real tract distributions were evaluated with a likelihood. Each population has four estimated likelihoods corresponding to the four Tracts models evaluated. A single run of each model was considered for each Mexican state, the one with the best likelihood. Finally, likelihoods were adjusted using the Bayesian Information Criterion (BIC), to account for the additional degrees of freedom of the more complex multi-pulse models versus the less flexible single pulse ones (**SI Table 1**). All best-fitting models consisted of an initial admixture event between Native Americans and Europeans followed by an African pulse. Admixture timing results are provided in generations and converted to dates by setting the date of the sampling to 2010 and assuming generations of 30 years, the average of the generation span of both genders [45].

### (f) Asian admixture timing in Guerrero with Tracts

Tracts only considers models of two and three ancestries. In order to estimate an admixture timing of the Asian component in Guerrero, the five continental ancestries from RFMix were merged into two categories, Asian and Non-Asian, since the only timing being estimated at this step was the introgression of the Asian ancestry into the admixed populations of Mexico. The East Asian and Melanesian assignment probabilities were combined into this “Asian” category, while sub-Saharan African, European and Native American components were merged into the general “Non-Asian” label. Three models were tested “pp”, “pc” and “cp”, where the first one represents a single admixture event between both ancestries and the rest involve one ancestry having a continuous pulse across several generations. The simplest single pulse model fit the empirical tract distribution best, yielding the highest likelihood (**SI Table 2**). Only individuals with >4% combined East Asian and Melanesian ancestry were included in the analysis.

### (g) Native American MAAS-MDS

For the two MAAS-MDS analyses, local ancestry was inferred with RFMix using five continental references separately by array. Each array was merged with whole genome sequencing data from 1000 genomes and HGDP in order to keep the highest number of SNPs for each of the five RFMix runs, to make more accurate local ancestry calls. The five continental references were identical to the panel used for global ancestry. East Asian and Melanesian components were considered together to localize the Asian ancestry origins. Local ancestry calls were performed with 789,054 SNPs for Array A, 518,409 SNPs for Array B, 794,029 SNPs for Array C, 722,489 SNPs for Array D and 356,143 SNPs from Array E (see Table 1).

Array A included both cosmopolitan Mexicans and Native Americans from MGDP. Each sample was genotyped with two arrays: Affymetrix 500 K and Illumina 550 K, resulting in a high number of SNPs. This array included seven cosmopolitan Mexican sampling locations from Sonora, Tamaulipas, Zacatecas, Guanajuato, Veracruz, Guerrero and Yucatan, as well as three Native Americans groups: Tepehuan, Northern Zapotec (from Ixtlan District, Northern Sierra in Oaxaca State) and Maya (from Campeche State). Array B included three cosmopolitan Mexican sampling locations from MGDP genotyped with the Illumina 550K array: Durango, Oaxaca and Campeche. Array C included eleven Native American groups from NMDP genotyped with the Affymetrix 6.0 array: Tarahumara, Huichol, Purepecha, Nahua (from Jalisco State), Nahua (from Highland Puebla), Totonac, Mazatec, Northern Zapotec (from Villa Alta District, Northern Sierra in Oaxaca State), Triqui, Tzotzil and Maya (from Quintana Roo State). Array D included Native Americans from three Nahua populations genotyped with the Axiom LAT 1 array: Nahua (from Xochimilco and San Pedro Atocpan, Mexico City), Nahua (from Necoxtla, Central Veracruz) and Nahua (from Zitlala, Central Guerrero). Array E included seven Native American groups genotyped with the Illumina 610-Quad and Illumina 650Y arrays: Pima, Yaqui, Purepecha, Mixtec, Southern Zapotec (from Sola de Vega District, Southern Sierra in Oaxaca State), Mixe and Maya (from Quintana Roo State).

The MAAS-MDS was applied to the Native American ancestry segments, that is, masking intercontinental components of sub-Saharan African, European, East Asian, and Melanesian origin, in both cosmopolitan Mexicans and indigenous individuals. The analysis was run using average pairwise genetic distances and only considering individuals with >10% Native American ancestry. Each of the ten cosmopolitan Mexican populations were merged into a single sample point for better clarity.

### (h) Asian MAAS-MDS with cosmopolitan Mexicans

The several RFMix runs per array with five continental references were performed just as with the previous native MAAS-MDS. East Asian and Melanesian components were combined. Local ancestry calls were performed with 803,636 SNPs for Array A, 518,409 SNPs for Array B, 561,340 SNPs for Array C, and 542,879 SNPs for Array D (see Table 1).

For this run we applied a >5% combined East Asian and Melanesian threshold to be considered in the MDS. Array A and Array B were identical to the cosmopolitan Mexicans from the Native MAAS-MDS. Array A included 7 cosmopolitan Mexican populations and Array B included 3 populations. After applying the combined East Asian and Melanesian ancestry threshold filter, only one individual from Sonora, one from Yucatan, twelve from Guerrero and one from Oaxaca were considered in the run. Array C included twelve population categories spanning East Asian, Southeast Asian (mainland and maritime) and Oceania genotyped with Affymetrix 6.0: Japan, Northern China, Southern China, Vietnam, Mindanao (Manobo), Negrito from Mindanao, Sumatra (Semende and Besemah), Borneo, Lesser Sunda Islands (Alor, Flores, Roti and Timor), Maluku Islands (Hiri and Ternate), Fiji and Papua New Guinea highlands. Array D included three Filipino sampling locations from [46] re-genotyped with Illumina OmniExpress Bead Chips in [47]: Igorot, Luzon and Visayas. To this array dataset was added two whole genome Igorot individuals from the Simons Genome Diversity Project [48].

The MAAS-MDS analysis considered the tracts from the combined East Asian and Melanesian ancestry merge, thus masking intercontinental components such as sub-Saharan African, European, and Native American, especially in cosmopolitan Mexicans. The analysis was run with average pairwise distances as a dissimilarity measure and only considering individuals with >10% Native American ancestry. All individual (averaging both haplotypes to create genotype dosage vectors) are plotted as a single point.

### (i) Ethics and Data accessibility

This work was conducted using publicly available data obtained through the respective Data Access and Material Transfer Agreements with the Institutions that published the data. Ethical procedures are thus described therein and access to the data and reference panels described here should be sought through the original sources as detailed in the References section.

## Authors’ contributions

AME and AGI conceived the study, designed the methodological approach, and supervised the project. AME, KS, RPE, CQC, and JERR selected samples and datasets. JERR, AGI, ELCH, and JBP analyzed the data. JERR, AGI and AME wrote the manuscript. All authors read and approved the manuscript.

## Acknowledgments

We thank the participants of the various studies assembled and reanalyzed here. We are grateful to Prof. Leonor Buentello (deceased, formerly at IIA-UNAM) for pioneering some of the community engagement and sampling efforts that derived in the integration of Nahua populations in reference panels. We thank Mitzi Flores and Adriana Garmendia for project management support and to Jacob Cervantes for IT support. We also thank Scott Huntsman, Celeste Eng, and Esteban G. Burchard for access to previously genotyped samples and genotype data curation, and Devang Agrawal for assistance with MDS analyses.

## Supplementary material

### Supplementary Figures

**Fig. S1:**
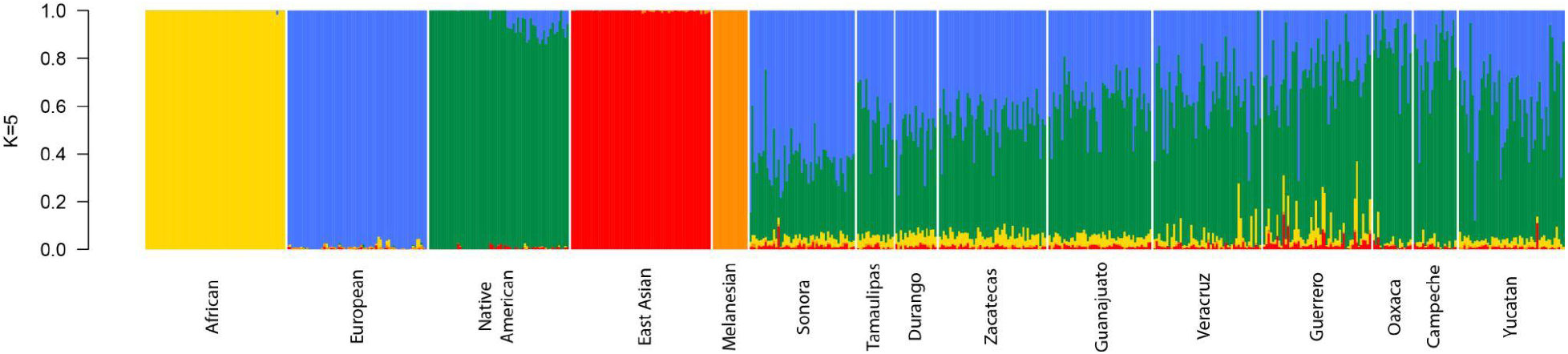
Admixture run at K=5 with 10 cosmopolitan Mexican and 5 continental reference populations.

**Fig. S2:**
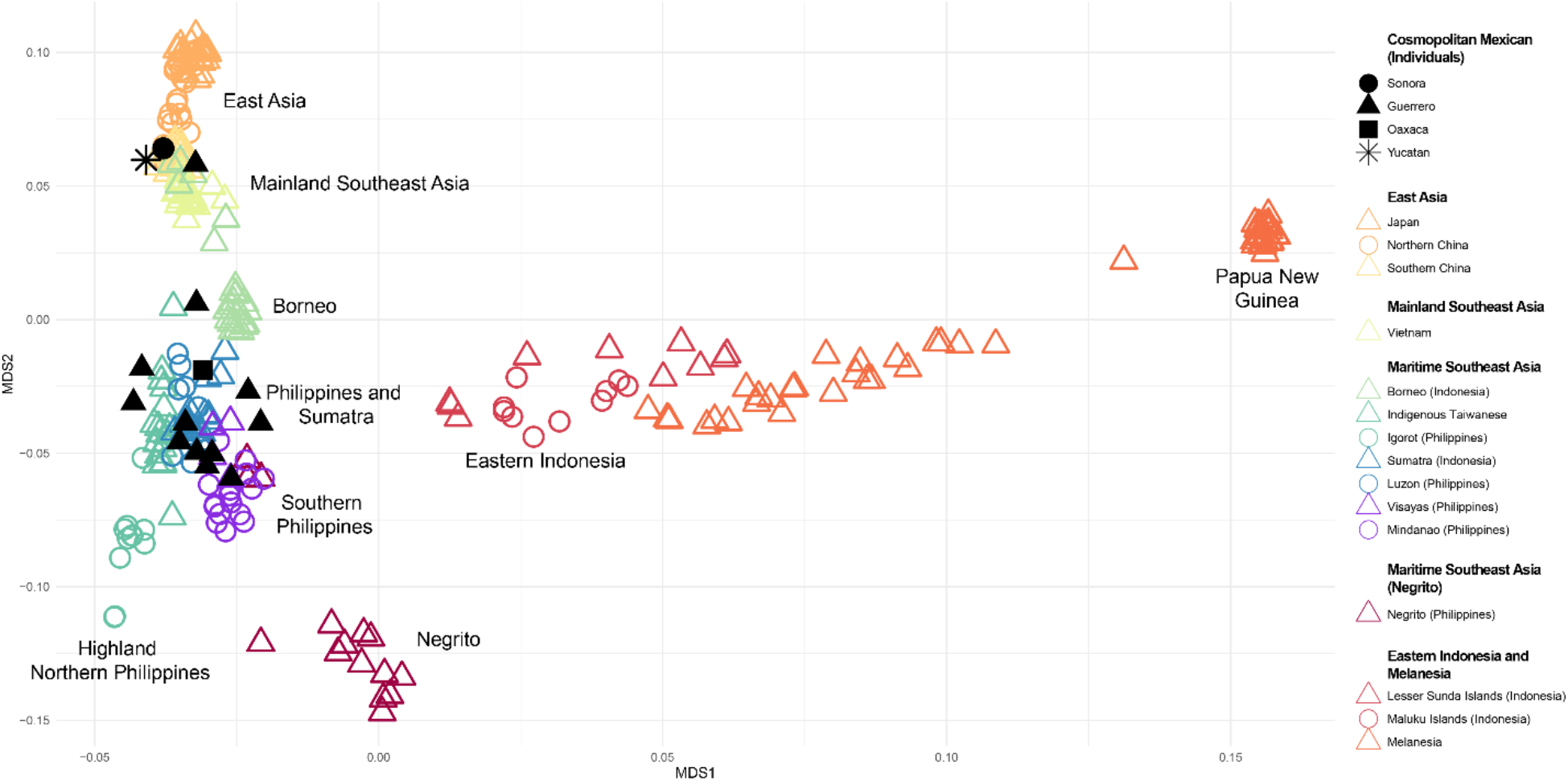
East and Southeast Asian MDS. Complete version of **Figure 3**, showing Melanesian variation on Papua New Guinea and eastern Indonesia on the right. Cosmopolitan samples did not exhibit affinity to these populations; therefore, MDS axis 1 was cropped in **Figure 3** to better visualize substructure in Mexico.

**Fig. S3:**
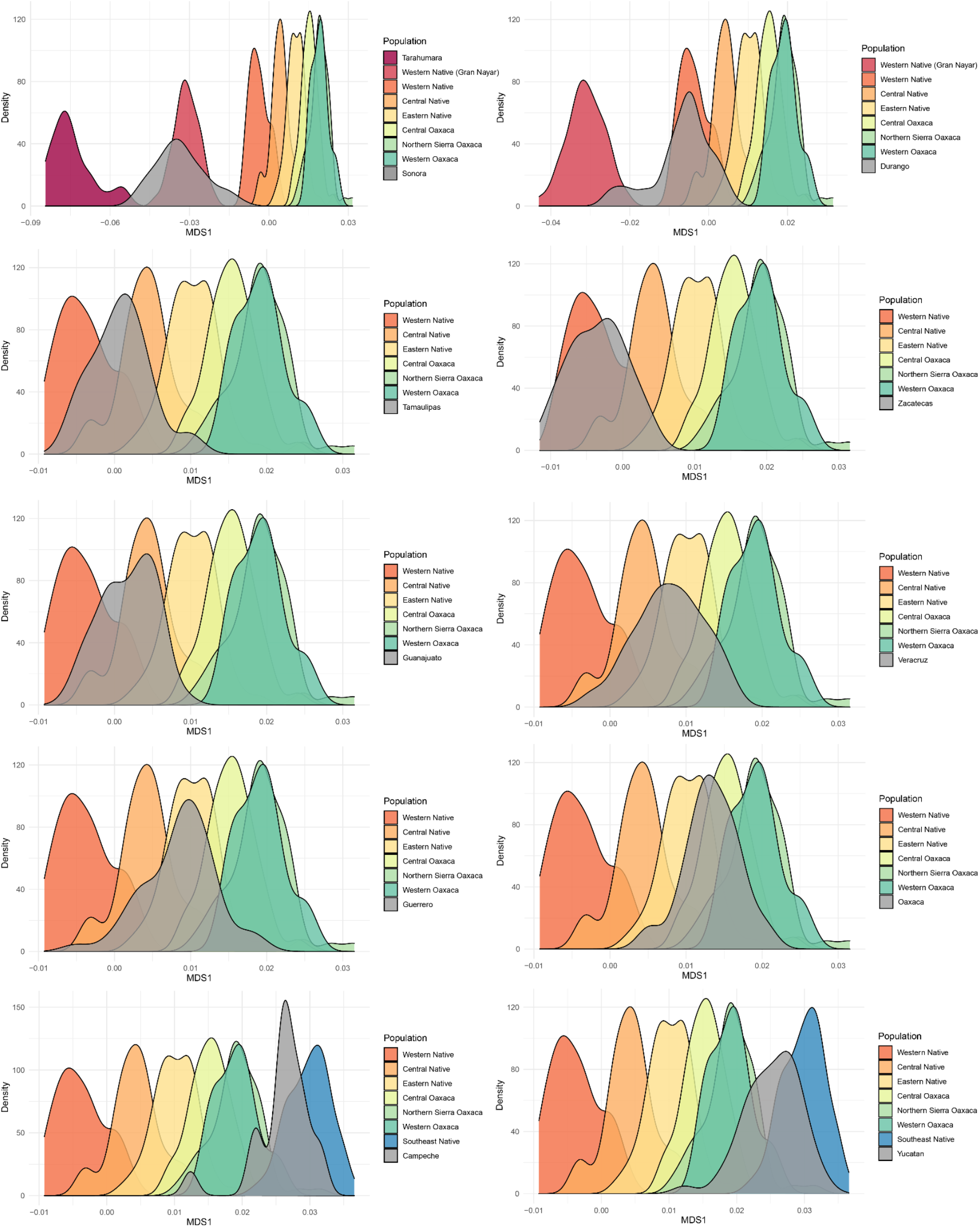
Native American MDS kernel density plot of 10 cosmopolitan Mexican populations. Each cosmopolitan population is plotted in a separate panel as a gray density distribution. This distribution considers all individuals instead of a combined data point from a population. The X axis corresponds to the MDS axis 1 from **Figure 2**.

### Supplementary Tables

**Table S1:**
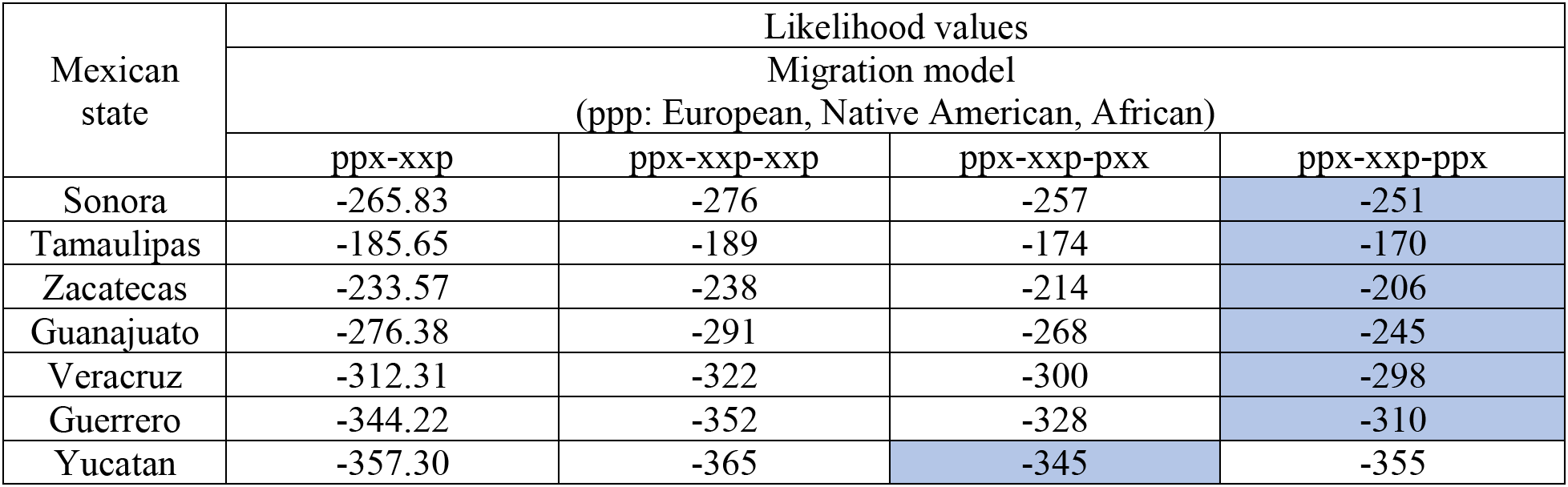
Admixture timing models from Tracts in cosmopolitan Mexicans at K=3. Four Tracts models were tested for each population, the models with the lowest likelihood (shown in blue) have the best fit to the real data. These models in blue correspond to the models shown in **Figure 1** and the histograms from **SI Figure 2**. Each model has an associated admixture pulse timing, as well as an expected histogram per ancestry that can be compared to the empirical ancestry histogram from the dataset. Most states show an initial European and Native American admixture event (ppx), followed by a single African pulse (xxp) and additional dual European and Native American pulses into the admixed population (ppx). Some states vary in their third admixture event, exhibiting a simple European pulse (Durango and Yucatan) or no third admixture event at all (Oaxaca and Campeche).

**Table S2:** Asian admixture timing models in Guerrero with Tracts. The model with the best fit to the real data, consists of a single discrete pulse between Asian and non-Asian ancestries 13 generations in the past. The timing of the admixture event is shown in parenthesis using the sampling date of 2010 CE and a 30-year per generation interval.

| Mexican state | Migration model |  |
| --- | --- | --- |
|  | (pp: Asian, Non-Asian) |  |
|  | pp<br>Single admixture pulse | pc<br>Multiple non-Asian pulses |
| Guerrero<br>(>4% of East Asian + Melanesian ancestry) | -131.17<br>(13 gen or 1620 CE) | -150.93<br>(10 gen or 1710 CE) |

